# First record of highly invasive Chinese sleeper *Perccottus glenii* Dybowski, 1877 (Perciformes: Odontobutidae) in the Elbe River Basin, Czechia

**DOI:** 10.1101/2023.01.20.524995

**Authors:** Marek Šmejkal, Ondřej Dočkal, Kiran Thomas, Lukáš Kalous

## Abstract

The Chinese sleeper (*Perccottus glenii*) has invaded European freshwaters and we present evidence of its first documented occurrence in the Elbe River basin in Czechia. The individual fish was caught by a fisherman and posted on social media. After immediately contacting the person in question, we obtained a live fish from him. The Chinese sleeper appears to have been present in interconnected ponds and streams for ten years and may have spread over a larger area. We recommend that eradication measures be implemented to prevent further spread.

**Statement of Significance:** The Chinese sleeper has invaded many European countries in the last 50 years, and it is expected to invade western Europe because of the favourable conditions for its establishment. This finding indicates that it could spread in the Elbe River Basin, which could have serious impacts on floodplain and wetland ecosystems. To prevent this scenario, eradication measures should be implemented.

## Brief communication

Introduced invasive alien fish species (IAS) are one of the greatest threats to freshwater diversity, affecting ecosystems through novel interactions of predation, competition, or parasite/disease transmission (Reshetnikov, 2004; Tapkir *et al*., 2022). The Chinese sleeper (*Perccottus glenii* Dybowski, 1877) is a significant threat to wetlands and ponds in Europe and is currently listed among the 26 priority IAS of animals in the European Union (European Commission, 2017). The Chinese sleeper occurs naturally in eastern Eurasia, ranging from the Amur Basin in Russia through north-eastern China to the north of the Korean Peninsula (Berg, 1949; Elovenko, 1981).

The Chinese sleeper was introduced from its original range into various European freshwaters (Reshetnikov, 2010). The history of its invasion in Europe starts with its release from aquaria in Saint Petersburg and Moscow in 1916 and 1940, respectively (Reshetnikov, 2004). Subsequently, in the last 50 years, it has spread mostly due to human interventions in Belarus (Lukina, 2011), Ukraine (Kvach, 2012), Lithuania (Reshetnikov, 2010), Poland (Grabowska *et al*., 2009), Latvia (Tambets & Järvekülg, 2005), Hungary (Harka, 1998), Slovakia (Koščo *et al*., 2003), Romania (Nalbant *et al*., 2004), Serbia (Gergely & Tucakov, 2003), Bulgaria (Jurajda *et al*., 2006), Estonia (Tambets & Järvekülg, 2005), Moldova (Mosu, 2007), Croatia (Ćaleta *et al*., 2011), Germany (Nehring & Steinhof, 2015; Reshetnikov & Schliewen, 2013), and Finland (Pihlström *et al*., 2022), and the summary of its invasion can be found in Reshetnikov (2010). As the Chinese sleeper has been detected in three out of four neighbouring countries of the Czech Republic, a high-risk potential for this country was assumed.

Chinese sleeper may become a dominant species in stagnant waters of floodplains and lentic waters with macrophytes and has serious impacts on insects, fish, and amphibians (Reshetnikov, 2001, 2003). Considerable declines were recorded in insect diversity due to Chinese sleeper predation on dragonflies and beetle larvae or adult aquatic beetles (Reshetnikov, 2003). Its impact on amphibians is alarming, as some species of newts (*Triturus cristatus, Lissotriton vulgaris*) and frogs (*Rana* sp., *Pelophylax lessonae*) have been reported to disappear from the invaded sites. In addition, their feeding niche overlaps with the European mudminnow (*Umbra krameri*), crucian carp (*Carassius carassius*) and sunbleak (*Leucaspius delineatus*), already threatened species with very similar preferred habitat (Glińska-Lewczuk *et al*., 2016; Koščo *et al*., 2008; Tapkir *et al*., 2022), resulting in interspecific competition between native species and the Chinese sleeper. Another described interaction between the Chinese sleeper and native fishes is predation on developing fish eggs and smaller size classes of fish (Reshetnikov, 2001, 2008), which can effectively eliminate other fish species in small water bodies. Therefore, any introduction of this IAS into previously pristine areas should be monitored with great awareness, and eradication campaigns are recommended to control its spread (Nehring & Steinhof, 2015; Pihlström *et al*., 2022).

For the Chinese sleeper, there are generally three typical pathways for human-caused introduction, which include recreational angling, the ornamental fish trade, and aquaculture. The beginning of its invasion history in Europe is associated with aquarium fish breeding in Moscow (Reshetnikov, 2004). However, the further spread is more likely due to stocking for aquaculture, with the first major event occurring in Ukraine (Reshetnikov, 2010, 2013). Another pathway, as with other highly invasive fishes, as it is the transport of bait fish for angling (Kalous *et al*., 2013; Reshetnikov, 2013). Finally, source populations may naturally spread downstream, especially during elevated floodplain river levels where surrounding unconnected pools serve as source populations for the colonization of downstream areas (Reshetnikov, 2013).

Because of the difficulty of monitoring freshwaters and the high cost compared to surveys in terrestrial environments, harnessing available sources of information collected from the public can help monitor biodiversity with particular attention to IAS in freshwater (Jarić *et al*., 2020a). Jarić et al. (2021) state that images posted on social media are important sources for detecting IAS and that images can be used to identify species and detect new introductions or range expansions. For example, the use of culturomics (human interaction with nature), citizen science (use of data collected by citizens for research/conservation), or iEcology (use of internet data generated for other purposes to study nature) can help reveal advances in the range of IAS (Jarić *et al*., 2020b; Ladle *et al*., 2016). Because angling is a widespread outdoor activity, especially in the western world, posting on social media or direct involvement of anglers in IAS monitoring can help track and suppress IAS (Arlinghaus *et al*., 2019; Cooke *et al*., 2016).

The present note describes the finding of a Chinese sleeper in the Czech Republic within the Elbe River Basin. The picture of a fish appeared in the Facebook group “<u>Ryby, které mizí z našich vod. Karas obecný, Slunka obecná a další naše …</u>” (Fishes disappearing from our waters. Crucian carp, sunbleak and other ours…) on January 10, 2023, together with the question whether it is “hlaváčkovec glenův”, which is the Czech name of the Chinese sleeper (Figure 1a). We contacted the Facebook user to confirm the location, date and other circumstances of the discovery and to get permission to share his photo for publication purposes. The person kept the fish alive until it was picked up by authors on January 13, 2023, and brought to the laboratory of the Institute of Hydrobiology, Biology Centre of the Czech Academy of Sciences in České Budějovice. There, the life of the specimen was sacrificed by an overdose of anaesthetics (MS −222) and morphometry (standard length, total length in mm, weight in g) and meristic counts were performed according to Horvatić et al. (2022) following the guide for first records (Bello *et al*., 2014). A picture was taken (Figure 1b) and the specimen was fixed in 40% formaldehyde solution and stored in the depository of the Institute of Hydrobiology.

**Figure 1:**
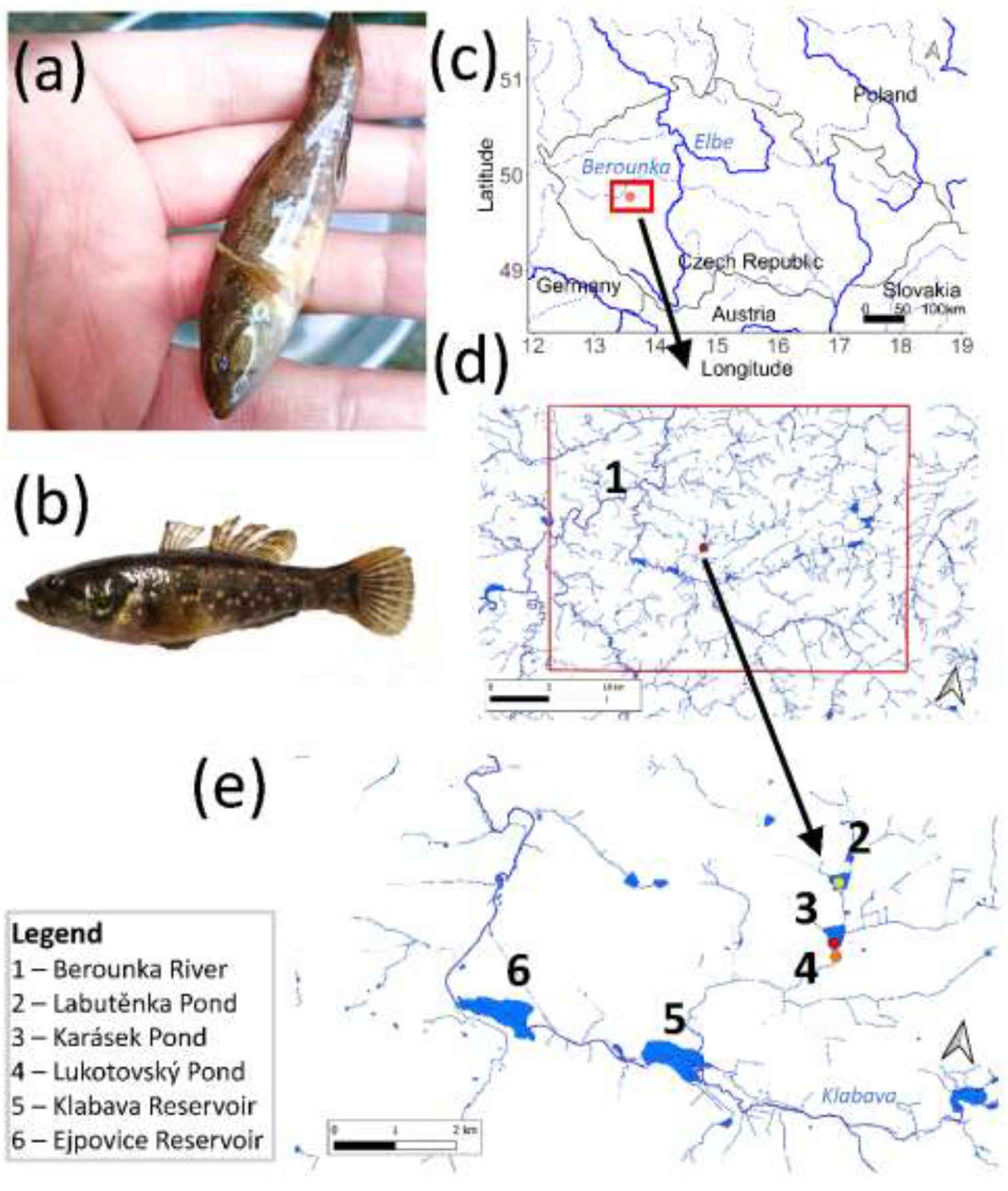
First record of the Chinese sleeper (*Perccottus glenii*) in the Elbe River Basin and the Czech Republic: (a) - the picture posted on social media, (b) - the picture of the specimen taken in the laboratory, (c) – the position of the record in the Elbe River Basin, (d) – the proximity of the pond system to the Berounka River, (e) – the system of three ponds interconnected with stream and surrounding larger water bodies. Yellow point – initial fish import according to manager, red point – current record, orange point – confirmed presence of the Chinese sleeper by pond manager. Maps were drawn using the program R version 4.1.1, the package ggmap (Kahle & Wickham, 2013; R Core Team, 2022) and the qGIS (QGIS Development Team, 2018) using layers of the State Administration of Land Surveying and Cadastre (CUZK, 2023).

On January 13, 2023, we contacted the company managing the pond to obtain more information on the possible origin of the Chinese sleeper and other related facts about its presence in the fish stock. In addition, we analysed the potential spread pathways downstream from the map and contrasted them with the information obtained from the company, as emptying the pond is an important source of Chinese sleeper to streams downstream (Nehring & Steinhof, 2015).

The fish comes from Karásek Pond in Osek u Rokycan municipality (N 49° 46’ 20” E 13° 35’ 17”), altitude 392 m.a.s.l. The pond with an area of 8.2 ha is situated in western Bohemia on the Osecký stream, a third-order tributary of the Berounka River, Elbe River Basin (Figure 1c-e). The fish was collected on January 8, 2023, during fish harvest in the stream below the dam. The morphometric and meristic characteristics were consistent with the species delimitation according to Horvatić *et al*. (2022). The specimen had a standard length of 67 mm, a total length of 79 mm, and a weight of 8.1 g. Eight meristic features on the left side of the body matched the species description: six rays in the first dorsal fin, 13 rays in the second dorsal fin, 13 rays in the pectoral fin, 4 rays in the ventral fin, 9 rays in the anal fin, 38 scales in the lateral line, 14 scales in the transverse row, 9 scales in the circumpenducular row (Horvatić *et al*., 2022; Nikolić *et al*., 2021). To our knowledge, the Chinese sleeper has not yet been detected in the Czech Republic (Chobot & Němec, 2017; Froese & Pauly, 2000); however, its range includes almost all neighbouring countries, Germany, Poland, and Slovakia, except for Austria (Grabowska *et al*., 2009; Koščo *et al*., 2003; Reshetnikov & Schliewen, 2013) and thus its invasion was expected.

According to the fish farm manager, the species identified in this work as Chinese sleeper occurs sporadically in the Labutěnka, Karásek and Lukotovský ponds (Figure 1e). The fish has been present in these three ponds for at least ten years. The pond manager linked the occurrence of the Chinese sleeper to the stocking of the Labutěnka pond with imported juvenile fish from Hungary. It seems very likely that the Chinese sleeper was present as contamination of the imported stock since it has been present in Hungary since 1997 (Harka, 1998).

The risk analysis of a non-native species is of considerable value in terms of analysing the potential negative impacts (ecological, economic, social) it may have on the invaded ecosystem and region (Verreycken, 2015). Strong dispersal potential has been predicted in areas of central and western Europe, high climatic suitability, and higher potential for human-mediated transfer (pond aquaculture) place these regions at higher risk of invasion (Rechulicz *et al*., 2015; Verreycken, 2015). The Chinese sleeper is defined as an ecologically plastic species that has a high establishment capacity and can tolerate significant fluctuations in water level, temperature, and dissolved oxygen (Bogutskaya & Naseka, 2002). The broad diet of the Chinese sleeper indicates that it is a non-selective, opportunistic predator (Grabowska *et al*., 2009; Reshetnikov, 2013). This highly flexible feeding strategy favours the spread of Chinese sleeper in invaded waters (Grabowska *et al*., 2009). Considering its invasion history, it has spread remarkably fast over a very large area.

Chinese sleeper is among the 26 animals defined by the European Union as IAS of exceptional interest (European Commission, 2017), and is also among the 27 major IAS of animals introduced into Europe for aquaculture and other related activities (Savini *et al*., 2010). The introduction of other parasites for which the Chinese sleeper acts as a host poses an additional threat to its potential for disease transmission (namely *Gyrodactylus perccotti* (Monogenea) (Ondračková *et al*., 2012), the nematode parasite *Philometriodes parasiluri* (Nikolic *et al*., 2007) and the cestode *Nippotaenia mogurndae* (Kosuthova *et al*., 2004)).

Characteristics such as high resilience, broad habitat requirements, voracity, and ease of reproduction mean that the Chinese sleeper has significant impacts on species composition and especially on threatened species such as amphibians, aquatic insect species, and floodplain fishes (Rechulicz *et al*., 2015). Its resilience allows it to colonize habitats where other fish predators are unsuccessful due to the lack of oxygen, so the interconnected fauna of these ecosystems are likely not adapted to the change in predation.

Pond aquaculture has a tradition in the Czech Republic, with the most important fish species being the common carp (*Cyprinus carpio*); however, this could provide fertile ground for the Chinese sleeper invasion of Czech waters. Pond aquaculture also produces coarse fish species that are partially flushed from the pond during draining and harvesting, and among them are often small fish species such as topmouth gudgeon (*Pseudorasbora parva*) (Gozlan *et al*., 2010; Kajgrová *et al*., 2022; Musil *et al*., 2014). In addition to the release of Chinese sleeper from ponds into streams (Nehring & Steinhof, 2015), populations established in streams may be fed by this influx of small prey (Bojarski *et al*., 2022). Additionally, common carp from pond aquaculture is commonly stocked into fishing grounds in rivers, which emphasises the importance of fish stock control (Kalous *et al*., 2018). The landscapes filled with ponds and reservoirs are more susceptible to fish invasions (Šmejkal *et al*., 2023), as has also been shown by the spread of IAS topmouth gudgeon and gibel carp (*Carassius gibelio*) (Lusk *et al*., 2010).

As for the situation in the identified occurrence area, three kilometres in the southwestern direction downstream is the Klabava Reservoir with an area of 39.2 ha and rich riparian vegetation. The Chinese sleeper prefers waters that either have weak flow or are stagnant, with well-developed aquatic vegetation, floodplains with well-developed vegetation, riparian zones of lakes, marshy waters, and even swamps (Horvatić *et al*., 2022; Reshetnikov, 2004). Reaching this reservoir can support them with a suitable habitat for their establishment, where measures will be very difficult to implement.

The finding was immediately reported to the Ministry of the Environment of the Czech Republic. Although we do not currently know how far Chinese sleeper has spread from the identified area of occurrence, the finding initiated immediate steps to monitor and eradicate Chinese sleeper from the area to prevent it from spreading further downstream in the Elbe River watershed.

## Acknowledgements

We thank Roman for reporting this important record on the Facebook group and raising awareness of this threat to biodiversity.

## Ethical statements

### Ethical approval

The field sampling methods and experimental protocols used in this study were performed by the guidelines and permission from the Ministry of Environment of the Czech Republic (OZPZ/2022/IN/1).

### CRediT authorship contribution statement

**Marek Šmejkal**: Conceptualization, Writing- Original draft preparation, Reviewing and Editing, Funding acquisition. **Ondřej Dočkal:** Writing- Original draft preparation, Visualization. **Kiran Thomas**: Data curation, Writing- Original draft preparation. **Lukáš Kalous:** Conceptualization, Methodology, Writing- Original draft preparation, Reviewing and Editing, Funding acquisition.

